# Loss of mu and delta opioid receptors on neurons expressing dopamine receptor D1 has no effect on reward sensitivity

**DOI:** 10.1101/2020.03.18.996454

**Authors:** Zofia Harda, Jadwiga Spyrka, Kamila Jastrzębska, Łukasz Szumiec, Anna Bryksa, Marta Klimczak, Maria Polaszek, Sławomir Gołda, Joanna Zajdel, Anna Błasiak, Jan Rodriguez Parkitna

## Abstract

Opioid signaling controls the activity of the brain’s reward system. It is involved in signaling the hedonic effects of rewards and also has essential roles in reinforcement and motivational processes. Here, we focused on opioid signaling through mu and delta receptors on dopaminoceptive neurons and evaluated the role these receptors play in reward-driven behaviors. We generated a genetically modified mouse with selective double knockdown of mu and delta opioid receptors in neurons expressing dopamine receptor D1. Selective expression of the transgene was confirmed using immunostaining. Knockdown was validated by measuring the effects of selective opioid receptor agonists on neuronal membrane currents using whole-cell patch clamp recordings. We found that in the nucleus accumbens of control mice, the majority of dopamine receptor D1-expressing neurons were sensitive to a mu or delta opioid agonist. In mutant mice, the response to the delta receptor agonist was blocked, while the effects of the mu agonist were strongly attenuated. Behaviorally, the mice had no obvious impairments. The mutation did not affect sensitivity to the rewarding effects of morphine injections or social contact and had no effect on preference for sweet taste. Knockdown had a moderate effect on motor activity in some of the tests performed, but this effect did not reach statistical significance. Thus, we found that knocking down mu and delta receptors on dopamine receptor D1-expressing cells does not appreciably affect reward-driven behaviors.

**Highlights:**

– It is well accepted that opioid signaling controls the brain’s reward system
– We generated mutant mice with mu and delta receptor knockdown in D1 neurons
– Knockdown made dopaminoceptive neurons insensitive to mu and delta opioid receptor agonists
– The mutation did not cause obvious behavioral impairments
– The loss of mu and delta receptors on D1 neurons does not affect reward sensitivity

## 1. Introduction

Opioid signaling controls the activity of the brain’s reward system, acting as a regulator of the mesolimbic system, both at the level of dopamine neurons in the midbrain and dopaminoceptive medium spiny neurons in the striatum. It is now well established that mu receptor agonists inhibit GABAergic signaling in the ventral midbrain, which leads to an increase in dopamine neuron activity and is critical for the euphoric and reinforcing effects of opioids (Fields and Margolis, 2015). Conversely, the role of opioid signaling in dopaminoceptive neurons (i.e., neurons that receive dopamine inputs), particularly striatal medium spiny neurons, remains only partly understood. It has been reported that the activation of mu or delta opioid receptors in discrete areas of the nucleus accumbens of the striatum induces hedonic reactions and promotes the intake of palatable food or drink (e.g. Castro and Berridge, 2014; Peciña and Berridge, 2005). These observations led to an influential model linking opioid signaling through mu and delta receptors in the nucleus accumbens to the “liking” component of reward, and separating it from dopamine signaling, involved primarily in motivational processes, or the “wanting” component of reward (Berridge et al., 2009; Wise, 2004). Nevertheless, some evidence has indicated that such elegant separation of roles may be too simplistic. The reported reinforcing effects of direct mu opioid receptor agonist injection into the striatum on conditioned place preference in rats appear to be inconsistent, although anatomical differences have been suggested as a potential explanation for these discrepancies (Bals-Kubik et al., 1993; Castro and Berridge, 2014). Intrastriatal injections of delta agonists have been observed to have no effect on place preference (Bals-Kubik et al., 1993). The complete loss of mu opioid receptors abolishes the rewarding effects of morphine in mice (Matthes et al., 1996), and the reintroduction of mu receptor expression to direct pathway medium spiny neurons of the striatum is sufficient to restore morphine-conditioned place preference (Cui et al., 2014). It has also been reported that the selective deletion of mu opioid receptors in forebrain GABAergic neurons (including those in the striatum) spares opioid-conditioned place preference but affects motivation to self-administer heroin or obtain palatable food (Charbogne et al., 2017). To add further complexity, it appears that opioid signaling in the forebrain may play different roles depending on the type of reward, as the same mutation completely abolishes alcohol-conditioned place preference (Hamida et al., 2019). It should also be noted that mu opioid signaling in the nucleus accumbens has been implicated in the rewarding effects of social interaction (e.g. Trezza et al., 2011). Conversely, data regarding the specific roles of delta opioid receptors in reward signaling are more limited. A recent report implicated these receptors in resilience to social stress through a mechanism involving primarily D1-expressing neurons (Nam et al., 2019). Thus, while the majority of reports have implicated opioid signaling in the striatum in the control of reward-driven behaviors, there is no consensus on the actual mechanisms involved.

All types of opioid receptors are expressed in the striatum (including the nucleus accumbens), and their mRNA and protein levels follow discrete distribution patterns (Le Merrer et al., 2009; Mansour et al., 1995; Svingos et al., 1996). Mu opioid receptor mRNA and protein are mainly present and ligand binding mainly occurs in striatal patches or striosomes (Crittenden and Graybiel, 2011). Conversely, delta opioid receptor mRNA levels are relatively low, but the delta opioid receptor is the most abundant opioid receptor protein in the striatum. It has been reported that delta opioid receptors are present mainly on indirect pathway neurons that project to the intermediate nuclei in the basal ganglia while mu receptors are expressed both on direct and direct pathway medium spiny neurons; however, the reported fractions of neurons expressing these receptors vary considerably (Ambrose et al., 2006; Banghart et al., 2015; Oude Ophuis et al., 2014). Recent single-cell transcriptome analyses of the whole brain identified a subclass of neurons that express dopamine receptor D1 but have no or very low delta opioid receptor expression as well as subtypes of cells that express the D1, mu and delta receptors (Zeisel et al., 2018). Together, these data suggest that the expression of opioid receptors in the same neuronal population may be variable, and there is no clear agreement among reports based on different methodologies.

Here, we investigated the role of mu and delta opioid receptor-dependent signaling in D1-expressing neurons in reward-driven behaviors. We generated a novel mouse model with selective knockdown of both receptors by means of RNA interference. The main rationale for targeting both receptors was that they are mainly activated by the same ligands – enkephalins – and trigger intracellular signaling cascades initiated by G_o/i_ proteins (Gendron et al., 2016; Williams et al., 2013). Therefore, while these receptors may potentially have different roles in the striatum, this is mainly due to their presence on different cell types; on the other hand, their effects in a single type of neuron are highly similar. Thus the elimination of one of these receptors could be compensated by the effects of the other. We found that while knockdown reduced or abolished the activity of the mu and delta receptors, no significant effects on reward-driven behaviors were observed.

## 2. Materials and Methods

### 2.1. Animals

#### 2.1.1. Oprd1/Oprm1^D1-KD^ mouse strain

Short hairpin RNAs against the mu and delta receptors were designed using BLOCK-IT RNAi Designer software (Invitrogen, USA). The sequences of the 21-nt fragments complementary to the target mRNAs were GCTGCCCTTTCAGAGTGTTAA (Oprm1-1), CCTTTGGAAACATCCTCTGCA (Oprm1-2), GCTGGTGATTCCTAAACTGTA (Oprd1-1) and CGCCTTGAGATAACATCGGGT (Oprd1-2). Synthetic oligos were cloned into the GW/EmGFP-miR vector (Invitrogen), which contains sequences derived from miR-155 (see Fig. 1A for summary). Silencing efficiency was validated in CHO-K1 cells cotransfected with plasmids encoding opioid receptors and plasmids encoding the miRNA cassette. A construct harboring 4 hairpins was recombined into a bacterial artificial chromosome (BAC; RP24–179E13; Children’s Hospital Oakland Research Institute, USA) containing the mouse *Drd1a* gene. The BAC was purified, the vector sequences were removed, and the transgene was injected into the pronuclei of fertilized oocytes from SWR/J mice. Mutant mice were congenically bred onto the C57BL/6N background (>8 generations of backcrosses before the start of the experiments).

**Fig. 1.**
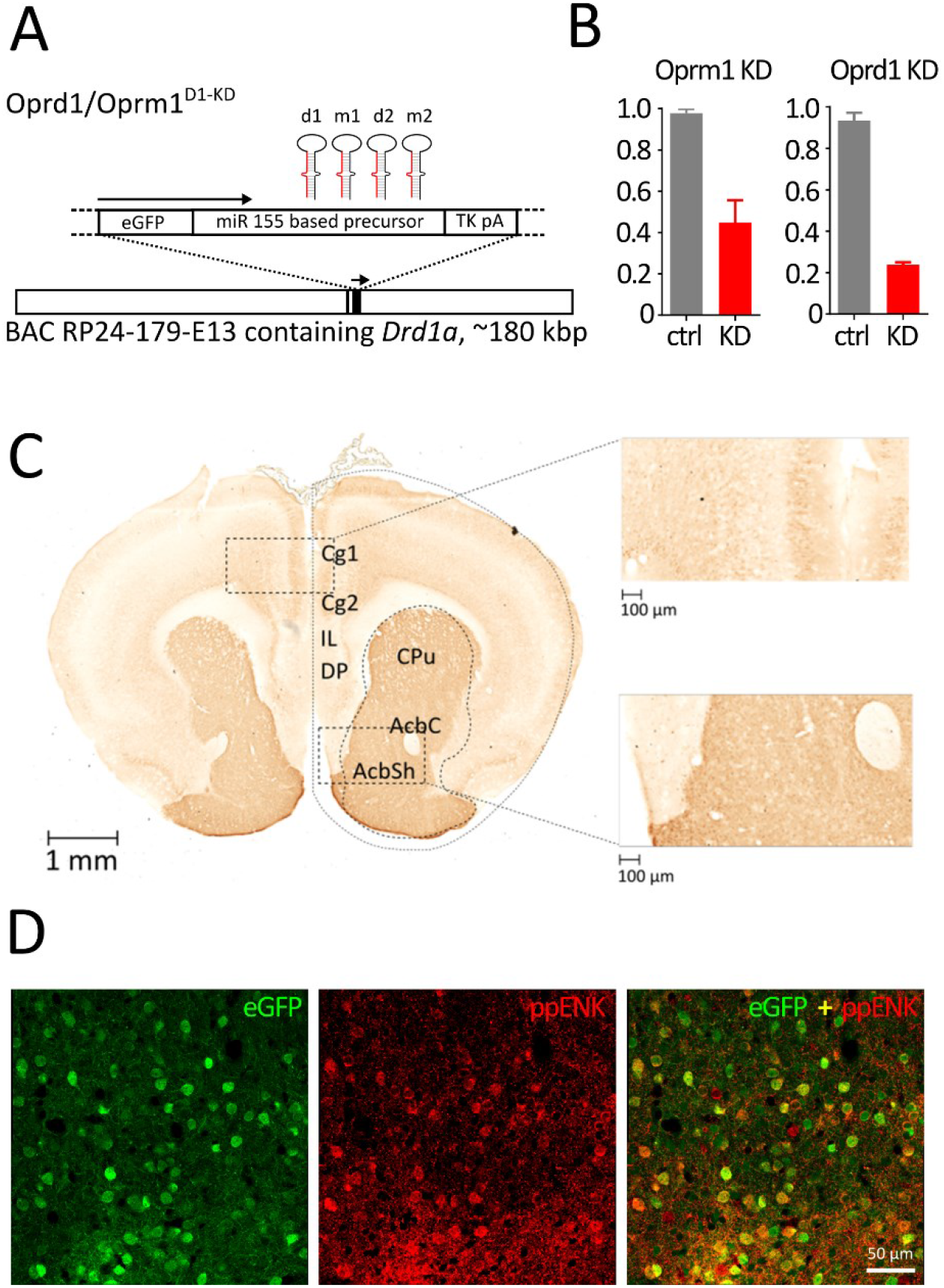
Generation of the Oprd1/Oprm1^D1-KD^ mouse strain. **(A)** Schematic representation of the construct used to generate the mouse strain. **(B)** Summary of knockdown efficiency results in CHO-K1 cells. The results show that the levels of Oprm1 or Oprd1 mRNA fragments were normalized after the cells were cotransfected with a construct expressing the targeted receptor fragment together with a construct expressing corresponding hairpins or a control hairpin. **(C)** The micrograph shows immunological staining of GFP encoded by the transgene. The images on the right are magnifications of the selected areas. Abbreviations: Cg – cingulate cortex, IL – infralimbic cortex, DP – dorsal peduncular cortex, AcbC – nucleus accumbens core, AcbSh – nucleus accumbens shell. **(D)** The micrographs of the nucleus accumbens show fluorescent immunostaining for GFP and preproenkephalin (ppEnk) and the composite image.

Transgenic animals were genotyped by PCR using primers with the sequences ACGTAAACGGCCACAAGTTC and ACGTAAACGGCCACAAGTTC (amplicon size 180 bp), which target the eGFP-encoding sequence. Additionally, each genotyping reaction also included positive control primers with the sequences CCATTTGCTGGAGTGACTCTG and TAAATCTGGCAAGCGAGACG (amplicon size 370 bp). The PCR conditions were as follows: 3 minutes of denaturation at 95°C followed by 40 cycles of 30 s at 95°C, 30 s at 58°C, and 60 s at 72°C.

The experiments were performed on adult *Oprd1/Oprm1*^D1-KD^ mice and their wild-type littermates aged 7-17 weeks at the beginning of the procedures. The exception was the socially conditioned place preference procedure, which was started when mice reached the age of 4 weeks. Male mice were used for all experiments except the IntelliCage experiment. The animals were housed 2-6 per cage in rooms with a controlled temperature of 22±2°C under a 12/12-h light-dark cycle with *ad libitum* access to food and water. The animals were handled for 5-7 days before the beginning of the experimental procedures. The experiments were conducted during the dark phase, except for the light-dark box test, and morphine-conditioned place preference test, which were conducted during the light phase. The saccharin preference test and the IntelliCage experiment were conducted over one or several days.

All behavioral procedures were approved by the II Local Bioethics Committee in Krakow (permit numbers 1000/2012, 224/2016, and 264/2019) and conducted in accordance with the European Communities Council Directive of 24 November 1986 (86/609/EEC).

#### 2.1.2. Transgene expression

The expression of the transgene was assessed by analyzing the coexpression of the eGFP protein with the miR precursor by immunostaining. Both immunohistochemistry and immunofluorescence were used. Briefly, the animals were perfused with 4% paraformaldehyde in phosphate-buffered saline, and the brains were removed and postfixed overnight. For immunochemistry, the brains were sliced into 40-μm sections in the coronal plane on a vibratome (Leica, Germany), the sections were blocked with goat/pig serum hybridized with an anti-GFP antibody (rabbit, Thermo Fisher Scientific, A-11122, 1:500), and then the signal was developed using the Rabbit Vectastain ABC HRP kit and diaminobenzidine substrate (Vector Inc., USA). Images were acquired using an Aperio ScanScope CS device (Leica, Germany).

For immunofluorescence, perfused brains were frozen in 30% (w/v) sacharose, sliced into 40-μm sections on a cryostat (Leica), and mounted on Superfrost+ microscope slides. The sections were stained with an anti-GFP antibody (chicken, Abcam, ab13970, 1:10000) and, in some cases, a rabbit anti-ppEnk antibody (Neuromics, RA14124-50; 1:1000). The following secondary antibodies were used: Alexa Fluor 488-conjugated goat anti-chicken IgY (H+L) (Invitrogen, A-11039, 1:1000) and Alexa Fluor 555-conjugated donkey anti-rabbit (Invitrogen, A-31572, 1:1000). Fluorescence images were acquired using an LSM700 Zeiss upright confocal microscope.

### 2.2. Whole-cell patch clamp activity recordings

#### 2.2.1 Brain slice preparation

Mice (6- to 10-week-old males) were decapitated under isoflurane (Aerrane, Baxter) anesthesia. Coronal brain slices (250 μm thick) containing the nucleus accumbens were cut on a vibratome (VT 1000S Leica Microsystems, Germany). The slices were prepared from brain tissue submerged in ice-cold cutting artificial cerebrospinal fluid (ACSF) containing (in mM) 92 NaCl, 30.0 NaHCO_3_, 1.25 NaH_2_PO_4_, 10.0 MgSO_4_, 2.5 KCl, 0.5 CaCl_2_, 20.0 HEPES, 5.0 sodium ascorbate, 3.0 sodium pyruvate, 2.0 thiourea and 10 glucose and continuously bubbled with a mixture of 95% O_2_ and 5% CO_2_, pH 7.3 – 7.4, osmolarity 290-300 mosmol/kg. After cutting, the slices were immediately transferred to an incubation chamber filled with normal ACSF containing (in mM) 124 NaCl, 26 NaHCO3, 1.25 NaH2PO4, 1 MgSO4, 4.5 KCl, 1.8 CaCl2 and 10 glucose at 32 ± 0.2°C. The tissue was incubated for at least 120 minutes before electrophysiological recordings.

#### 2.2.2. Cell visualization and cell identity confirmation

Nucleus accumbens neurons were visualized with an upright microscope (Zeiss Axio Examiner A1 microscope, Zeiss, Germany) using video-enhanced infrared differential interference contrast (DIC) and fluorescence optics. Individual neurons were visualized using a 40× water immersion lens. Dopamine receptor D1-expressing cells were identified based on the expression of GFP (KD animals) or tdTomato (control animals). GFP was excited at 455 nm and tdTomato was excited at 530 nm by LED illumination (Zeiss Colibri).

#### 2.2.3 Whole-cell patch-clamp recordings

Whole-cell current- and voltage-clamp recordings were made using the SEC-05X amplifier (NPI, Tamm, Germany). Signals were filtered at 3 kHz and digitized at 20 kHz using a Micro1401 converter (Cambridge Electronic Design (CED), Cambridge, UK) with Signal and Spike2 software (CED, UK). Patch micropipettes (7-9 MΏ) were pulled from borosilicate glass capillaries (Sutter Instrument, Novato, CA, USA.) using the Sutter Instrument P97 puller. The internal pipette solution contained (in mM) 125 K-gluconate, 20 KCl, 2 MgSO_4_, 10 HEPES, 4 Na2-ATP, 0.4 Na-GTP, 5 EGTA and 0.05% biocytin, osmolarity 290 – 300 mosmol/kg, pH 7.2 – 7.3. A liquid junction potential of +12.4 mV was calculated, and the data were corrected for this value.

To characterize the basic electrophysiological properties of D1 neurons, recordings were performed in normal ACSF. The membrane potential of the recorded neurons was held at −75 mV by continuous current injections, and voltage responses to rectangular current steps (500 ms in duration, −140 pA to +240 pA, 10-pA increments, with 5-s intervals between steps) and depolarizing current ramp (0.1 to 1 nA) were recorded. The membrane resistance, time constant and capacitance were measured from the voltage response to a −140 pA hyperpolarizing pulse. The excitability of the recorded neurons was examined using depolarizing current pulses from +10 pA to +240 pA, and for each cell, the number of action potentials was plotted against the intensity of the injected current. Action potential (AP) parameters, specifically the AP threshold, amplitude, 10-90 rise time, half width, AHP minimum and action potential peak time to AHP minimum time (AP peak to AHP), were measured for the first action potential evoked by the minimal depolarizing current step. The rheobase was defined as the minimal current necessary to induce an action potential and was determined using a current ramp protocol. Next, to characterize current-voltage relationships, cells were voltage-clamped at a command potential of −75 mV, and recordings were performed in the presence of 0.5 μM tetrodotoxin. Voltage steps from −120 mV to +10 mV (500 ms in duration, 10-mV increments) were delivered every 3 seconds, and steady-state current responses were measured.

To verify the presence of mu and delta opioid receptors on the membranes of D1-expressing neurons, cells were voltage-clamped at a command potential of −65 mV. All recordings were performed in normal ACSF containing tetrodotoxin (0.5 μM). After obtaining stable whole-cell current recordings for at least 15 minutes, the selective mu opioid receptor agonist DAMGO (1 μM) or the selective delta opioid receptor agonist DPDPE (1 μM) was added to ACSF-perfused slices. Recordings lasted for at least 25 minutes following batch application of the drugs.

#### 2.2.4. Statistical analysis

The values are given as the means ± SEM. In all experiments, p < 0.05 was considered significant. All data sets were tested for normal distribution, and outliers were excluded from the analysis (ROUT method, Q = 5%). The firing characteristics of the recorded neurons were analyzed by linear regression. Statistical significance between groups was determined using either Student’s t test or the Mann-Whitney test when applicable. Data analysis was performed using GraphPad Prism for Windows. A recorded neuron was classified as responsive to a drug if the whole-cell current of the neuron after drug application differed from the baseline by more than three standard deviations.

### 2.3. Behavioral procedures

Behavior was recorded using a Basler (ac1300 – 60 gm) camera and EthoVision 11.5 software (Noldus). The experimenter was blinded to the genotypes of the animals tested.

#### 2.3.1. Open field test

Walking initiation and open field exploration were measured during the adaptation period before the social interaction test. The mice were introduced to a novel transparent plastic cage (55 × 37.5 cm × 20.5 cm high) containing 0.5 cm of bedding. The latency to leave a 17×17-cm square (outlined digitally) was used as a measure of walking initiation. Latencies were measured manually from video recordings with BORIS software (Friard and Gamba, 2016). Leaving the square was defined as putting all four paws outside the square. The distance moved during the 30-minute habituation period was measured by EthoVision 11.5 software (Noldus, The Netherlands).

#### 2.3.2. Light-dark box test

The test was performed in a two-chamber apparatus (each chamber was 18 × 16 × 20 cm high). The light compartment was brightly lit (430 lux), and the dark compartment was dimly lit (20 lux). The animals were placed in the dark compartment and allowed to freely explore the apparatus for 5 minutes.

#### 2.3.3. Social interaction test

Partners (9- to 10-week-old male C57BL/6N mice, Charles River Laboratories, Germany) were habituated to our laboratory facilities for at least one week prior to the experiments. Focal animals were habituated to the test cage for 30 minutes before the test (55 × 37.5 × 20.5 cm, containing 0.5 cm of bedding). Immediately after habituation, the partner animal was placed in the cage, and the mice were allowed to interact for 10 minutes. Control animals were exposed to a novel object instead of a novel mouse. Aspen gnawing blocks (4 × 4 × 5 cm) were used as novel objects. The blocks were identical in shape and size to the blocks in the home cages of the mice but differed in smell. Time in proximity was measured from video recordings with EthoVision 11.5 software (Noldus, The Netherlands). Proximity was defined as < 8 cm between body centers.

#### 2.3.4. Saccharin preference in the home cage

Saccharin preference was assessed as described previously (Jastrzębska et al., 2016). Briefly, individually housed animals had 24-h access to two 25-ml graduated drinking bottles. One bottle was filled with water, and the other bottle was filled with 0.1% saccharin solution. Food was provided *ad libitum* on the cage floor.

#### 2.3.5. Saccharin preference with delay discounting

The test was performed in an IntelliCage (New Behavior, Switzerland), which allows for long-term monitoring of up to 15 mice living in a group with minimal interference from the experimenters. Mouse behavior was monitored using RFID chips (UNO PICO ID, AnimaLab, Poznań, Poland). The cage consisted of a housing area and four operant chambers (situated in the corners of the cage) equipped with sensors. The operant chambers (later referred to as “corners”) were accessible by only one animal at a time. Each of the corners allowed access to two bottles through a guillotine door. The experiment consisted of two phases: the adaptation phase (4 days) and the test phase (30 days). During adaptation, the mice had free access to water in all corners. During the test, the mice were provided 0.1% saccharin solution in two of the corners. The guillotine door closed 10 seconds from the first lick or immediately after the mouse left the corner. The delay from the moment the mouse was detected in the corner until the doors opened increased from 0.5 to 35 seconds every 48 h (an initial delay of 0.5 s lasted for 96 h). The positions of the saccharin and water bottles were exchanged every 24 h.

#### 2.3.6. Morphine-conditioned place preference

The test was conducted in automatic conditioned place preference (CPP) cages with three compartments (ENV-256C, Med Associates Inc., USA). One of the side compartments had black walls and a white floor, while the other had white walls and a black floor. On the first day of the procedure, the mice were habituated to the apparatus for 5 minutes (the floor was covered with white paper). On the second day, a pretest was conducted; the mice were placed in the central compartment and allowed to freely explore the whole apparatus for 20 minutes. On conditioning days, the mice received an i.p. injection of either morphine hydrochloride (10 mg/kg every other day) or saline and were immediately placed in one of the compartments for 40 minutes. The pairing of the compartment with morphine injection was biased, i.e., the mice were assigned to the compartment that was initially less preferred. The conditioning phase lasted eight days and was followed by a posttest, which was performed in the same manner as the pretest, on the ninth day.

#### 2.3.7. Social-conditioned place preference

The procedure was performed as previously described, with modifications (Dölen et al., 2013; Panksepp and Lahvis, 2007). Before the test, the animals were housed in groups of 2 to 6 on aspen shavings with aspen gnawing blocks (context A). The test consisted of three phases: the pretest, conditioning phase, and posttest (Fig. 7A). During the pretest, the animals were placed in a custom-made plastic cage (40×40×30 cm high) divided into two identical compartments by a transparent plastic wall with a 5×5-cm opening at the base. Each compartment contained a type of novel bedding (cellulose (Biofresh Performance Bedding, Absorption Corp, USA, 1/8’ pelleted cellulose), beech (P.P.H. “WO-JAR”, Poland, Trociny bukowe przesiane gat. 1), or spruce (LIGNOCEL^®^ FS 14, J. Rettenmaier and Sohne, Germany) and a gnawing block different from the one in the home cage in size and/or shape (contexts B and C). The mice were allowed to freely explore the cage for 30 minutes. The amount of time spent in each compartment was measured by EthoVision XT 11.5 software (Noldus, The Netherlands). After the pretest, the mice were returned to their home cages (context A). The next day, the mice were assigned to undergo social conditioning (housing with cage mates) for 24 h in one of the contexts used in the pretest followed by 24 h of isolate conditioning (single housing) on the other type of bedding. Two experiments were performed on separate groups of animals. In the first experiment, beech and spruce were used. Since all mice showed a preference for beech over spruce, spruce was chosen as the social bedding. In the second experiment, beech and cellulose were used. The mice showed no preference between these beddings, and the assignment of contexts was random (unbiased design). The results of the two experiments were pooled. Conditioning was repeated for 6 days (3 days in each context, alternating every day). The posttest was performed in the same manner as the pretest. The rewarding effects of the social context were measured by comparing the time spent in the social context during the posttest to the time spent in the social context during the pretest.

## 3. Results

### 3.1. Generation of *Oprd1/Oprm1*^D1-KD^ mice

The strain was generated following the general outline of the method described by (Novak et al., 2010). The transgene harbors a sequence encoding eGFP for easy detection of expression and two hairpins against each of the targeted sequences, the mu and delta opioid receptors (Fig. 1A). We first tested the knockdown efficiency in the CHO-K1 cell line and found average reductions of 60 and 80% in the abundance of the mRNA sequences corresponding to mu and delta receptors, respectively (Fig. 1B). Transgenic mice were generated by injecting the transgene construct (without the vector cassette) into fertilized oocytes. The offspring were screened for the presence of the transgene, and two founder lines were established. The line with higher transgene levels was selected for further experiments. The results indicated that the expression of GFP was consistent with the known pattern of dopamine receptor D1 protein expression (Fremeau et al., 1991) (Fig. 1C). The strongest signal was observed in areas corresponding to the striatum, including the nucleus accumbens and olfactory tubercle, and slightly weaker staining was also present in the deeper layers of the cortex and discrete areas of the septum. Importantly, in the nucleus accumbens, the expression of GFP and preproenkephalin exhibited minor overlap (Fig. 1D), which is again consistent with D1 receptors being mainly expressed on the neurons of the direct pathway equivalent in the nucleus accumbens (Gangarossa et al., 2013; Kupchik et al., 2015) (Fig. 1C & D).

### 3.2. Whole-cell patch clamp recordings of activity in response to opioid antagonist treatment

To characterize the influence of selective double knockdown of the mu and delta opioid receptors on the electrophysiological properties of D1 receptor-expressing neurons in the nucleus accumbens core, we used *Oprd1/Oprm1*^D1-KD^ [Tg/0] mice as the KD group and D1-tdTomato mice [Tg/0] as the control group. Only neurons expressing tdTomato (control group) or GFP (KD group) were recorded (Fig. 2A, B). In total, 32 neurons from the control and 26 from the KD group were included in the analysis. Whole-cell recordings revealed that mu and delta opioid receptor knockdown did not influence the shape of the action potentials of the examined neurons (Fig. 2C). Subsequent statistical analysis did not reveal any differences in the measured AP parameters (action potential threshold, amplitude, 10-90 rise time, half width, and AHP minimum and AP peak to AHP, Tab. 1). Double knockdown of the mu and delta opioid receptors also did not influence the membrane resistance, capacitance or time constant of the recorded neurons (Fig. 2D, E, F). The excitability of nucleus accumbens neurons was determined by testing the relationship between the firing rate and the intensity of the injected current (Fig. 2G). The firing characteristics of the recorded neurons were fitted by linear regression (Fig. 2H), and the parameters did not differ between the groups. The mean gain and the threshold current were not affected by mu and delta opioid receptor knockdown (gain: 145.6 ± 7.35 Hz/pA in the control group and 132.0 ± 8.93 Hz/pA in the KD group, t(39) = 1.187, p = 0.243; threshold current: 0.13 ± 0.008 nA in the control group and 0.11 ± 0.016 nA in the KD group; t(39) = 1.003, p = 0.322). Current-voltage relationships (I-V curves) were plotted using measurements of steady-state currents in response to a series of incremental voltage steps recorded in ACSF containing tetrodotoxin. Two-way repeated measures ANOVA revealed no significant differences in the current responses of KD and control neurons (main effect of treatment, F(1,56) = 0.469, p = 0.496; Fig. 2I).

**Fig. 2.**
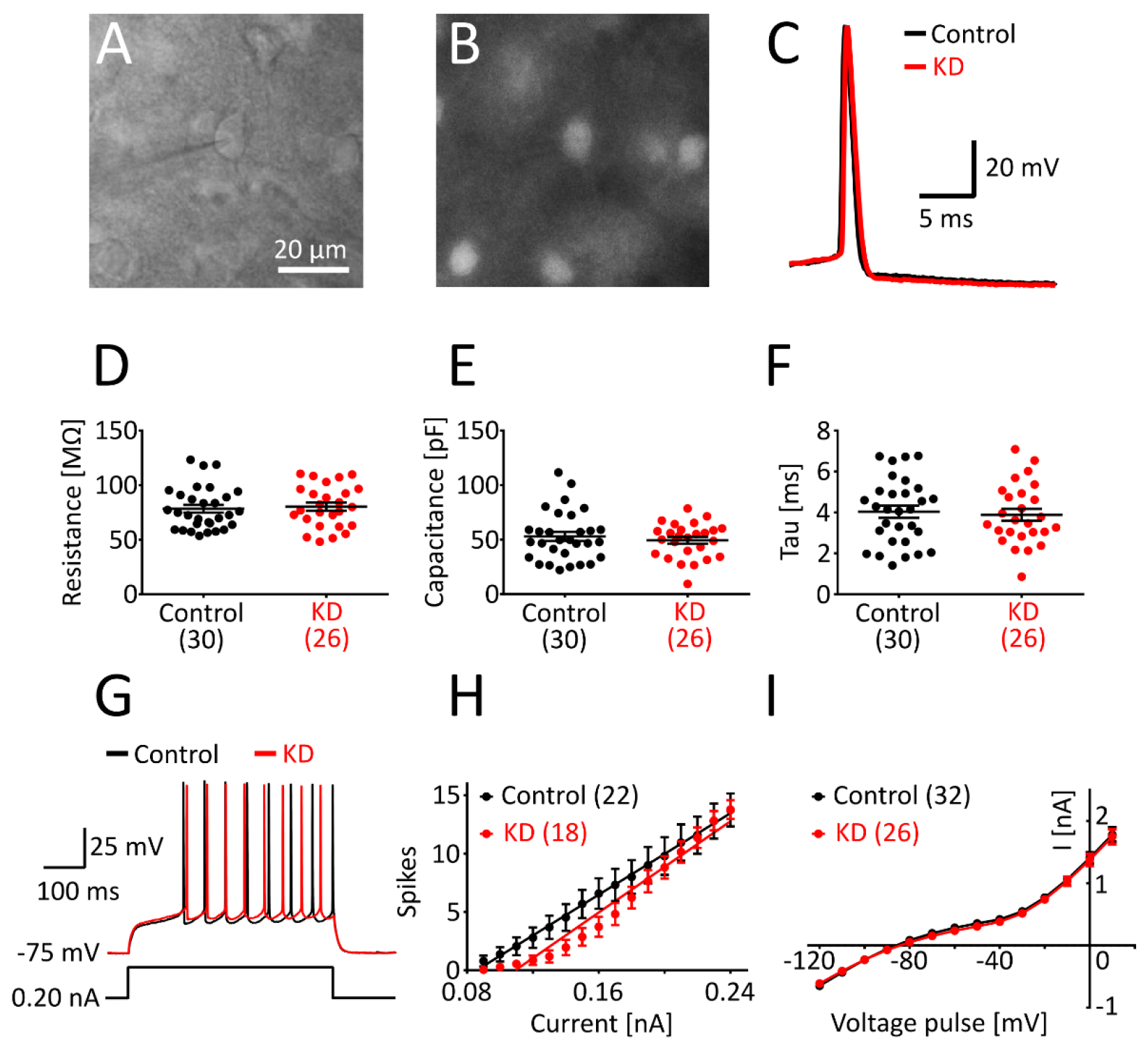
Passive and active membrane properties of nucleus accumbens core neurons expressing D1 receptor from control and Oprd1/Oprm1^D1-KD^ (KD) mice. (**A)** Infrared differential interference contrast (note the presence of the recording pipette) and **(B)** fluorescence images of nucleus accumbens core neurons expressing mCherry. **(C)** Typical current stimulus evoked action potentials recorded from neurons from control and KD mice. Comparison of **(D)** resistance, **(E)** capacitance, and **(F)** time constant (Tau) between neurons from control and KD mice. **(G)** Representative traces of membrane potential responses (upper traces) of control and KD neurons evoked by a 0.20-nA current step (lower trace). **(G)** Input-output relationship reflecting the excitability of neurons from control and KD mice. **(H)** Steady-state current-voltage relationship of neurons from control and KD mice recorded in the presence of tetrodotoxin (0.5 μM). The data are presented as the mean ± SEM, no statistically significant differences were revealed between examined neurons.

**Tab. 1.** Action potential parameters of nucleus accumbens core neurons expressing D1 receptor neurons from control and Oprd1/Oprm1^D1-KD^ (KD) mice

| Parameter | Control n=22 | KD n= 20 |
| --- | --- | --- |
| Threshold [mV] | -44.33 ± 0.64 | -43.75 ± 0.63 |
| 10-90 Rise [ms] | 0.33 ± 0.02 | 0.32 ± 0.01 |
| Amplitude [mV] | 73.28 ± 1.32 | 73.50 ± 1.52 |
| Half width [ms] | 1.09 ± 0.05 | 1.05 ± 0.04 |
| AHP minimum [mV] | -55.73 ± 0.60 | -57.02 ± 0.76 |
| Peak to AHP [ms] | 23.32 ± 1.74 | 23.61 ± 1.52 |

To validate the lack of functional mu opioid receptor expression in the D1-expressing neurons of *Oprd1/Oprm1*^D1-KD^ mice, we bath-applied the selective mu opioid receptor agonist DAMGO (1 μM) and recorded the whole-cell currents (–65 mV command potential) of D1-expressing cells in slices from KD and control mice. DAMGO induced a reversible outward current in 9 of 12 voltage-clamped control neurons (Fig. 3A, B). Statistical analysis of the DAMGO-induced current amplitude showed that in control cells, DAMGO application induced a statistically significant change in the recorded whole-cell current (mean whole-cell current at baseline: 0.23 ± 0.02 nA; mean whole-cell current during DAMGO application: 0.25 ± 0.02 nA; t(11) = 4.671, p = 0.0007; Fig. 3B). DAMGO application produced a reversible outward current in only 2 of 11 recorded KD neurons (Fig. 3C, D, E). Statistical analysis of the DAMGO-induced current amplitude showed that in KD neurons, DAMGO did not induce a statistically significant alteration in the whole cell current (mean whole-cell current at baseline: 0.23 ± 0.03 nA; mean whole-cell current during DAMGO application: 0.24 ± 0.03 nA; t(10) = 2.091, p = 0.063; Fig. 3D). Subsequent comparison of the DAMGO-induced whole-cell current in KD and control neurons showed that the mu opioid receptor agonist evoked a reduced effect in KD cells (whole-cell current change: 14 ± 6 pA) in comparison with control neurons (whole-cell current change: 28 ± 6 pA), but the difference did not reach statistical significance (t(21) = 1.686, p = 0.107; Fig. 3F).

**Fig. 3.**
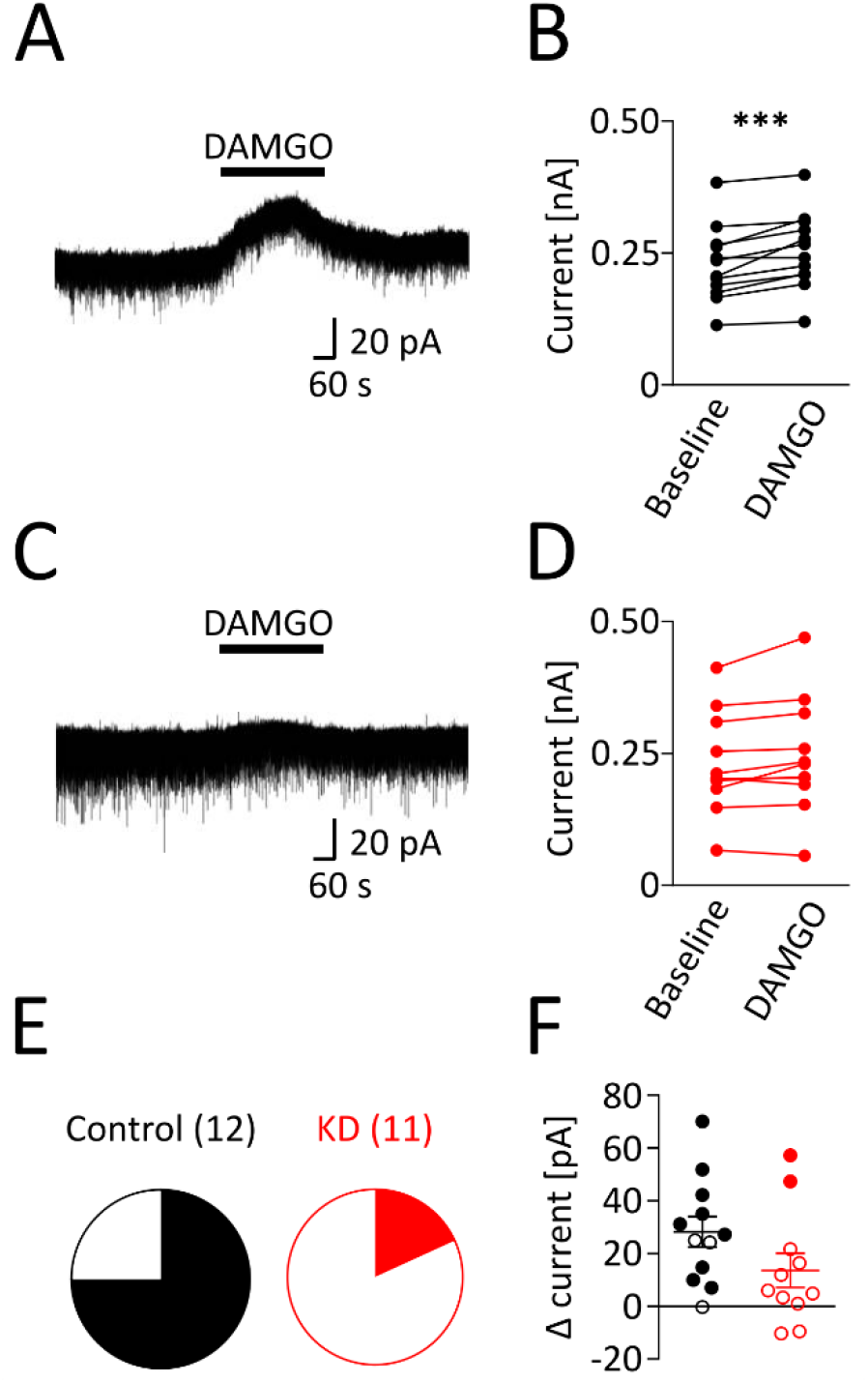
Influence of the delta opioid agonist DAMGO on the whole-cell current of D1 receptor-expressing striatal neurons. **(A)** A representative voltage-clamp recording showing DAMGO (1 μM, horizontal line)-induced outward whole-cell currents in control neurons and **(B)** a corresponding line graph showing the DAMGO-induced current amplitude. **(C)** A representative voltage-clamp recording showing that neurons from KD mice did not respond to DAMGO application and **(D)** a corresponding line graph showing the current amplitudes recorded at baseline and during DAMGO application. Note the lack of differences between whole-cell current amplitudes in the tested conditions. **(E)** Pie charts showing the proportion of DAMGO-responsive (filled) and DAMGO-nonresponsive (empty) neurons from control and KD mice. **(F)** The amplitudes of DAMGO-induced outward whole-cell currents recorded in neurons from control and KD mice. The filled circles represent neurons in which the change in current in response to DAMGO was greater than three standard deviations from baseline. The data are presented as the mean ± SEM. Significant differences between group means (Student’s t test) are represented by *** (p < 0.001).

To confirm the knockdown efficiency of delta opioid receptors in D1-expressing cells, D1 striatal neurons from *Oprd1/Oprm1*^D1-KD^ and control mice were voltage clamped at −65 mV, and their responsiveness to the selective delta opioid receptor agonist DPDPE (1 μM) was recorded (Fig. 4A, B, C, D). We found that all examined neurons from KD mice were insensitive to the delta opioid receptor agonist, which was confirmed by subsequent statistical analysis (mean whole-cell current at baseline: 0.26 ± 0.02 nA; mean whole-cell current during DPDPE application: 0.26 ± 0.02 nA; t(9) = 0.1545, p = 0.881). At the same time, in neurons from the control mice, bath application of the drug generated a reversible outward current (in 7 of 10 recorded cells, Fig. 4E), and the change in the whole-cell current was statistically significant (mean whole-cell current at baseline: 0.30 ± 0.02 nA; mean whole-cell current during DAMGO application: 0.31 ± 0.03 nA during DAMGO application; t(9) = 4.139, p = 0.0025). Moreover, the DPDPE-evoked change in the whole-cell current of D1 striatal neurons was significantly different between control and KD neurons (change in the whole-cell current in control neurons: 7 ± 2 pA; change in the whole-cell current in KD neurons: 0.2 ± 1.1 pA; t(18) = 3.411, p = 0.003; Fig. 4F).

**Fig. 4.**
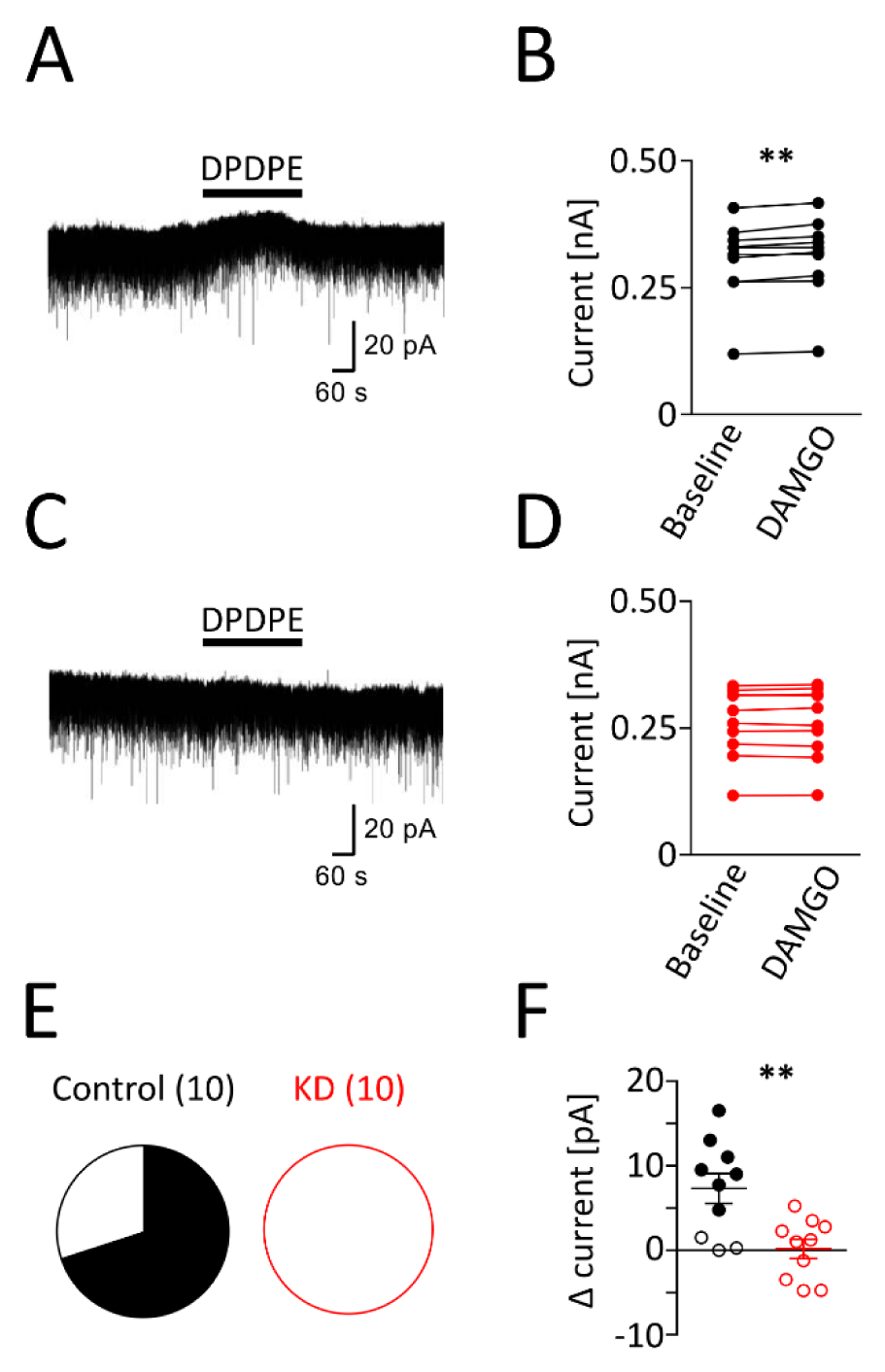
The influence of the mu opioid agonist DPDPE on the whole-cell current of D1 receptor-expressing striatal neurons. **(A)** A typical voltage-clamp recording showing DPDPE (1 μM, horizontal line)-induced outward whole-cell currents in control neuron and **(B)** a corresponding line graph showing the DPDPE-induced current amplitude. **(C)** A representative voltage-clamp recording showing that neurons from KD mice did not respond to DPDPE application and **(D)** a corresponding line graph showing the current amplitude recorded at baseline and during DPDPE application. Note the lack of differences between whole-cell current amplitudes in the tested conditions. **(E)** Pie charts showing the proportion of DPDPE-responsive (filled) and DPDPE-nonresponsive (empty) neurons from control and KD mice. **(F)** The amplitudes of DPDPE-induced outward whole-cell currents recorded in neurons from control and KD mice. The filled circles represent neurons in which the change in current in response to DPDPE was greater than three standard deviations from baseline. The data are presented as the mean ± SEM. Significant differences between group means (Student’s t test) are represented by ** (p < 0.01).

### 3.3. General phenotype of *Oprd1/Oprm1*^D1-KD^ mice

First, we assessed the behavior of mutant mice in an open field. The mutation targeted striatal neurons of the direct pathway, which may have an effect on motor activity. *Oprd1/Oprm1*^D1-KD^ mice showed a normal latency to leave the square in the open field test (Fig. 5A, t(40) = 0.562, p = 0.578), indicating no deficit in movement initiation. There was a trend towards a shorter total distance traveled by mutant mice compared to wild-type littermates, but this difference did not reach significance (t(40) = 1.7, p = 0.097; Fig. 5B). These results indicate no appreciable motor impairments that could confound the interpretation of the observed behaviors.

**Fig. 5.**
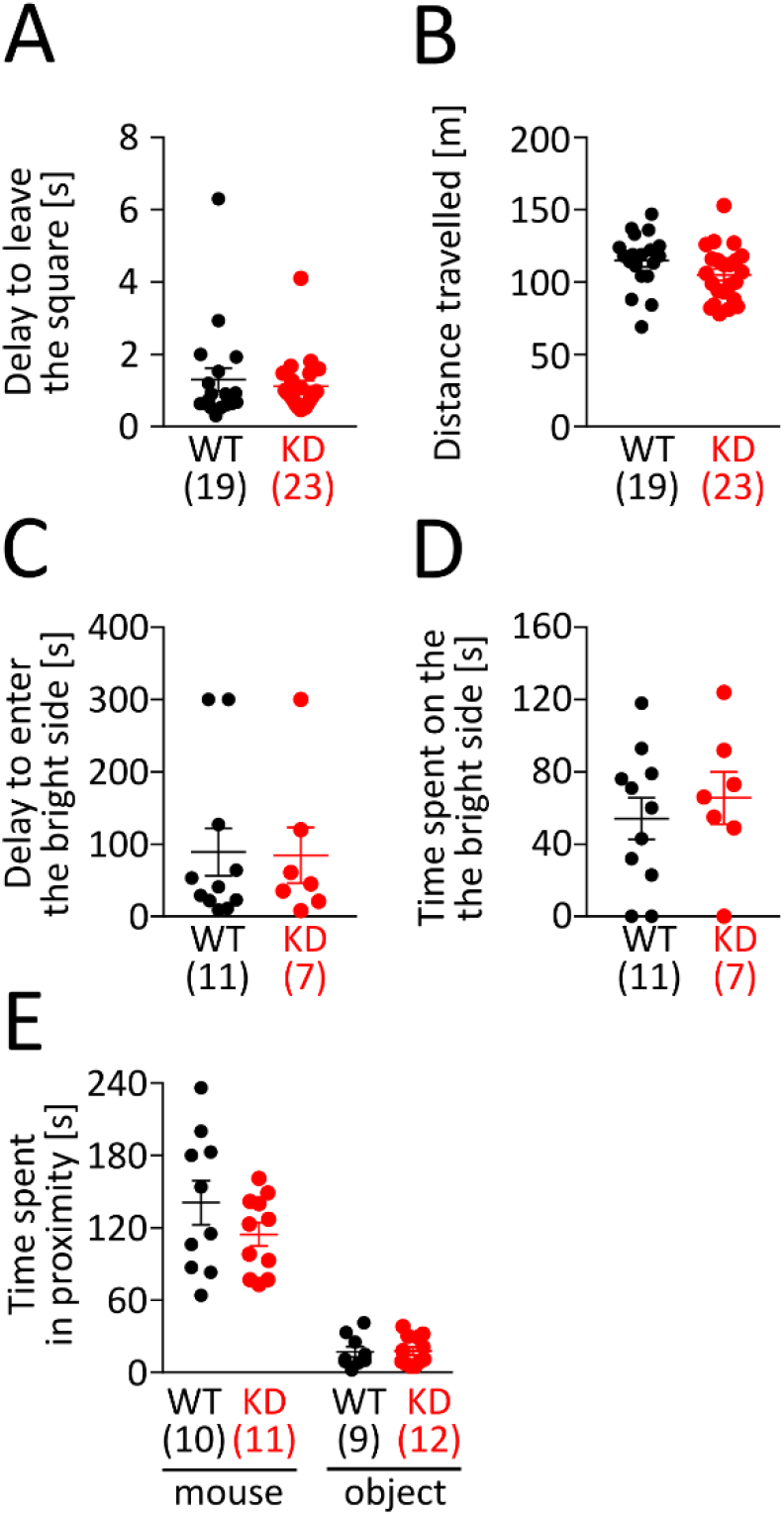
Locomotor activity, anxiety and social interactions in Oprd1/Oprm1^D1-KD^ mice. **(A)** Initiation of movement. The graph shows the latency for each mouse to leave a 17 × 17-cm square surrounding its initial position. **(B)** Total distance moved during the open field test. **(C)** Latency to enter the lit/bright compartment during the light-dark box test. A latency of 300 s indicates that the animal never left the dark part of the box. **(D)** Time spent in the lit/bright compartment. **(E)** Time spent in proximity of an unfamiliar conspecific or inanimate object during the interaction test (less than 5 cm between body centers). In all graphs, each point represents a single animal, while the bars and whiskers correspond to the means ± SEMs.

Next, we tested anxiety-like behaviors in the light-dark box and during social interaction tests. *Oprd1/Oprm1*^D1-KD^ mice did not differ from wild-type controls in the latency to enter the lit compartment (t(16) = 0.915, p = 0.928; Fig. 5C) and overall spent a similar amount of time on the lit side (t(16) = 0.622, p = 0.543; Fig. 5D). There was no effect of genotype on the number of crossings between compartments (control: 11.45 ± 2.31; *Oprd1/Oprm1*^D1-KD^: 11.71 ± 2.47; p = 0.945). We performed a social interaction test, another test for altered anxiety-like behavior, in the open field apparatus. *Oprd1/Oprm1*^D1-KD^ mice spent as much time in proximity with an unfamiliar adult conspecific as their wild-type littermates and spent significantly more time interacting with another mouse than an inanimate object (genotype: F(1, 38) = 1.487, p=0.23; animal vs. object: F(1, 38) = 110.8, p < 0.0001; animal vs. object × genotype: F(1, 38) = 1.66, p = 0.2015; Fig. 5E). Together, these results indicate that the mutation did not altered anxiety-like behaviors, indicating that its effects were different from the reported effects of the complete inactivation of the *Oprd1* gene (Filliol et al., 2000) or treatment with a delta opioid receptor agonist (Perrine et al., 2006).

### 3.4. Reward-conditioned behaviors in *Oprd1/Oprm1*^D1-KD^ mice

To assess the effects of the mutation on reward sensitivity, we used two paradigms: the saccharin preference test and morphine-or social-conditioned place preference. First, we measured the volume of saccharin-sweetened water (0.1% w/v) consumed by male *Oprd1/Oprm1*^D1-KD^ and wild-type mice over a period of 24 h in a two-bottle choice task. Mutant mice did not differ from wild-type animals in preference for sweet taste (t(8) = 1.479, p = 0.177; Fig. 6A) or the total volume of 0.1% saccharin solution consumed (t(8) = 0.118, p = 0,909; Fig. 6B). Next, we subjected female *Oprd1/Oprm1*^D1-KD^ and wild-type to a delay discounting task with saccharin as a reward. The main advantage of this method is that it is able to detect even a minor difference in subjective reward value and the propensity for impulsive choices. The test was conducted in IntelliCages, in which a group of 12 female mice implanted with radio-frequency identification (RFID) chips were housed together for 34 days. *Oprd1/Oprm1*^D1-KD^ mice showed similar general activity as control animals, which decreased slightly but significantly over the course of the experiment (number of corner visits, time: F(14, 140) = 5.783, p < 0.0001; genotype: F(14, 140) = 1.438, p = 0.258; time × genotype : F(14, 140) = 0.28, p = 0.995; Fig. 6C). The mutation had no effect on initial preference for the saccharin solution and did not affect the rate of discounting with increasing delay to access the reward (time: F(14, 140) = 99.75, p < 0 .0001; genotype: F (1, 10) =0.245, p = 0.636; time × genotype: F(14, 140) = 0.159, p = 0.9; Fig. 6D). There were no differences in the volume of water (time: F(14, 140) = 88.51, p < 0.0001; genotype: F(1, 10) = 0.005, p = 0.943; time × genotype: F(14, 140) = 0.190, p = 0.999; Fig. 6E) or saccharin solution (time: F(14, 140) = 94.72, p < 0.0001; genotype: F(1, 10) = 0.733, p = 0.412; time × genotype: F_(14, 140)_= 0.361, p=0.983; Fig. 6F) consumed. Thus, we found that the mutation did not affect preference for, the intake of or the motivation to drink the saccharin solution.

**Fig. 6.**
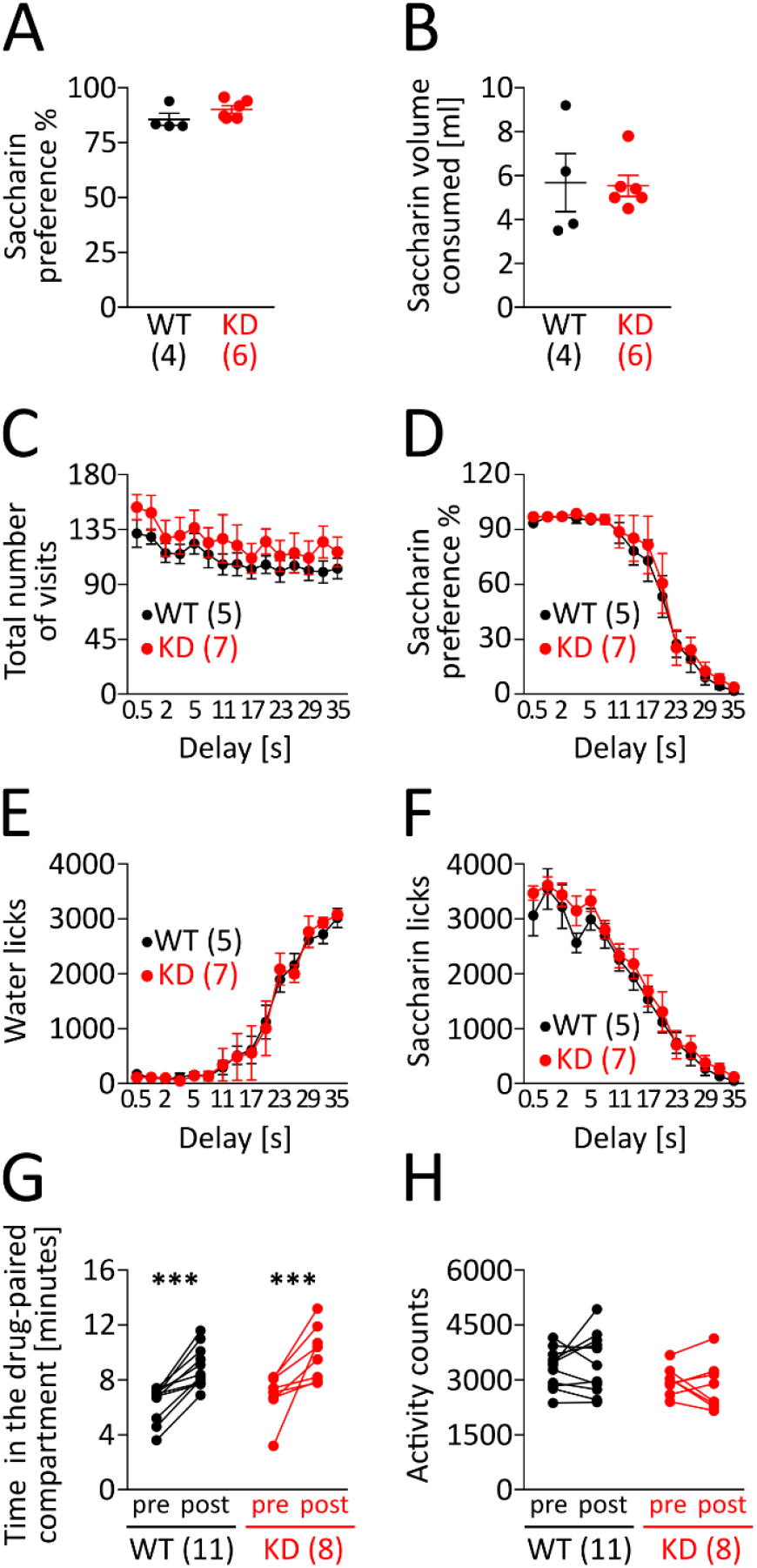
Reward sensitivity in Oprd1/Oprm1^D1-KD^ mice. **(A)** Preference for a 0.1% saccharin solution over water in the two-bottle test. **(B)** Total volume of the saccharin solution consumed over the duration of the test (24 h). The bars and whiskers correspond to the means ± SEMs **(C)** Activity in the IntelliCage during the delay-discounting test. The graph shows the mean number of visits per animal per 24-h period. The x axis shows the delay of access to reward. The error bars represent the SEMs**. (D)** Mean preference for a 0.1% saccharin solution over water during the delay-discounting procedure. **(E, F)** Mean number of licks of a water bottle and a saccharin bottle per 24 **(G)** Time spent in the morphine-paired compartment before (pre) and after (post) conditioning. Each pair of points connected by a line corresponds to an individual animal. **(H)** Motor activity in the apparatus before (pre) and after (post) conditioning. Activity was scored each time an animal crossed one of the infrared beams in the compartments. Significant differences between group means (Sidak’s test) are represented by *** (p < 0.001).

Next, we tested morphine-conditioned place preference (CPP). Both wild-type and *Oprd1/Oprm1*^D1-KD^ mice showed an increase in preference for the context paired with morphine injections (10 mg/kg, i.p.; pre-post: F(1, 17) = 45.41 p<0.0001; genotype: F(1, 17) = 1.748 p = 0.204; pre-post × genotype: F(1, 17) = 0.278 p = 0.605; post hoc Sidak’s test; wt: p = 0.0003; KD: p = 0.0004; Fig. 6G). Mutant mice had, on average, slightly lower locomotor activity (particularly during the posttest); however, this effect did not reach significance (pre-post: F(1, 17)= 0.0006, p = 0.989; genotype: F(1, 17) = 3.316, p = 0.0862; pre-post × genotype: F(1, 17) = 1.07, p = 0.316; Fig. 6H). The results showed no appreciable effects of receptor knockdown on morphine reward, which is consistent with previously reported normal CPP in animals with selective deletion of the mu receptor in forebrain GABAergic neurons (Charbogne et al., 2017).

Finally, we tested the rewarding effects of social contact using the conditioned place preference paradigm (Fig. 7A). Both mutant and control mice acquired a preference for the context associated with group housing (pre-post: F(1, 13) = 21.1, p = 0.0005; genotype: F(1, 13) = 1.54, p = 0.237; interaction: F(1, 13) = 0.195, p = 0.666; post hoc Sidak’s test; wt: p = 0.009; KD: p = 0.019; Fig. 7B). Motor activity was not affected by genotype, although animals from both groups exhibited an increases in activity from the pretest to the posttest (pre-post: F(1, 13) = 19.73, p = 0.0007; genotype: F(1, 13) = 2.9, p = 0.11; interaction: F(1, 13) = 0.054, p = 0.82; post hoc Sidak’s test; wt: p = 0.026; KD: p = 0.009; Fig. 7C). These results showed that the mutation did not appreciably affect the rewarding effects of social contact. We attribute the genotype-independent change in activity to the normal physical development of young animals.

**Fig. 7.**
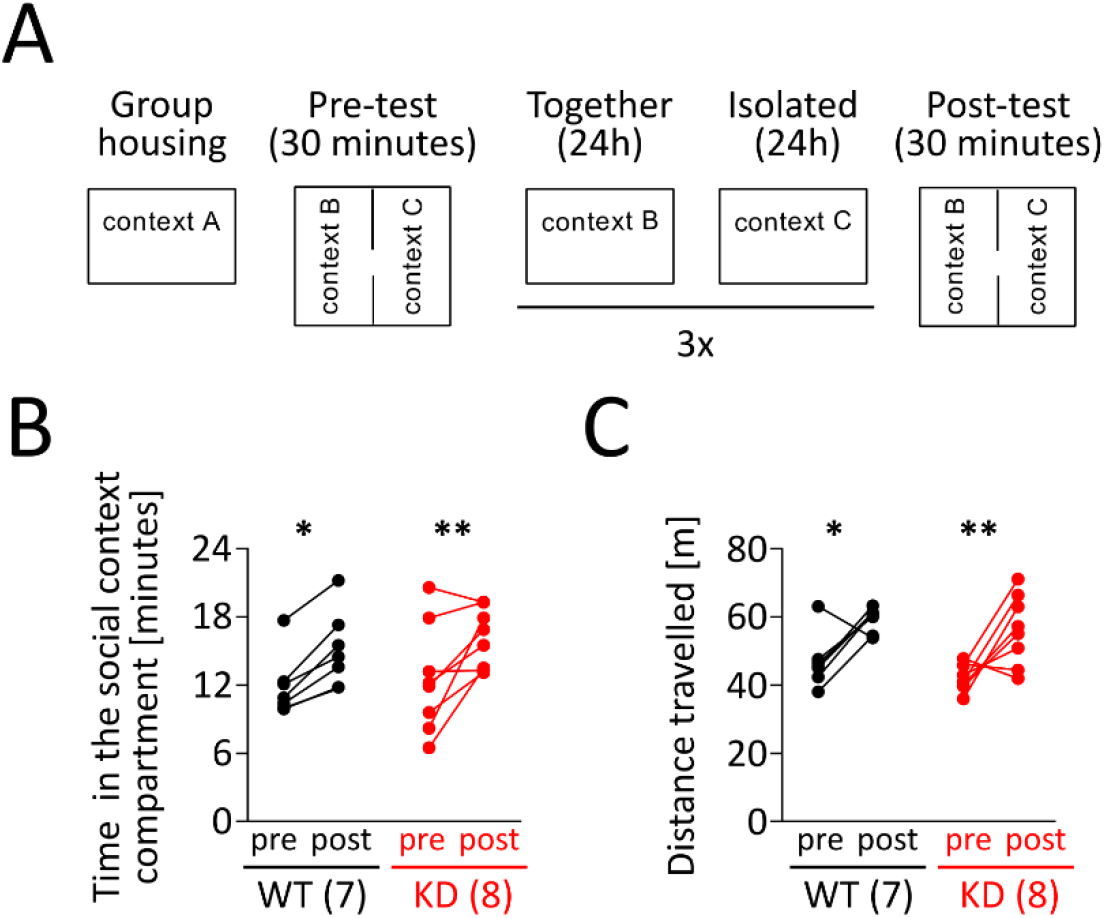
Rewarding effects of social interactions in Oprd1/Oprm1^D1-KD^ mice. **(A)** Schematic representation of the experiment. **(B)** Time spent in the context associated with social interaction before and after conditioning (in the pretest and posttest). Each pair of points connected by a line corresponds to an individual animal. **(C)** Total distance traveled during the pretest and posttest. Significant differences between group means (Sidak’s test) are represented by * (p < 0.05) and ** (p < 0.01).

## 4. Discussion

We found that knocking down the mu and delta opioid receptors in D1-expressing neurons had no appreciable effects on sensitivity to rewards or motivation to obtain them. The only changes in behavior were alterations in motor activity in some of the tasks that included rewards. These results do not completely exclude the involvement of the mu and delta opioid receptors present on neurons expressing dopamine receptor D1 in reward processing but suggests that they are not an essential part of the associated mechanism.

First, it should be stressed that the mutation utilized in this study targets neurons expressing dopamine receptor D1, which is a broader group of neurons than a subset of nucleus accumbens or striatal medium spiny neurons that prominently includes glutamatergic neurons in lower cortical layers. We note, however, that knockdown efficiency is dependent on gene promoter activity, and D1 expression is several-fold higher in medium spiny neurons than in other cell types (Nam et al., 2019). No significant behavioral effects of knockdown were observed, which could indicate that opioid signaling in non-striatal cells also has no essential role in reward-driven behaviors. However, it is likely that knockdown in cells other than medium spiny neurons was less efficient and that mu and delta opioid receptor signaling was thus spared. We also note that the knockdown efficiency in D1-expressing medium spiny neurons was probably higher than that found in the in vitro experiment based on the previously reported effects of mGluR5 knockdown in a mutant mouse line generated using the same gene promoter (Novak et al., 2010).

We showed that under *ex vivo* conditions, the majority of D1-expressing striatal medium spiny neurons from control mice were sensitive to mu and delta receptor agonists. The coexpression of the D1 and delta opioid receptors has been previously reported in striatal neurons (Ambrose et al., 2006; Ma et al., 2012). Here, we confirmed these data and showed that delta opioid receptor stimulation in D1-expressing neurons activated outward whole-cell currents, which correspond to the hyperpolarization of the cell membrane. To the best of our knowledge, the presented results are the first to show the direct, inhibitory effect of opioid receptor activation on whole-cell currents in striatal neurons. Notably, in knockdown mice, the mutation led to the complete loss of sensitivity to the delta opioid receptor agonist (DPDPE). These results not only confirm knockdown but also, along with results obtained in ACSF in the presence of tetrodotoxin, are proof of the presence of delta opioid receptors on postsynaptic membrane of the majority of striatal D1-receptor-expressing neurons. Similarly, most of the tested control cells were sensitive to the mu opioid receptor agonist (DAMGO), and the selective activation of mu receptors also led to outward current activation. D1 and mu opioid receptor coexpression in the striatum is well documented (Cui et al., 2014; Oude Ophuis et al., 2014), and its hyperpolarizing influence is attributed to postsynaptic action (Elghaba and Bracci, 2017). Again, the lack of response of knockdown mice to the agonists allows us to conclude that the functional mu opioid receptor is largely absent in mutant animals. Importantly, the lack of either mu or delta receptors did not change the passive or active membrane properties of D1-expressing neurons; therefore, any behavioral effect of the mutation can be attributed to altered opioid signaling, not the dysfunction of D1-expressing neurons.

The normal preference for sweet taste and morphine- or social-conditioned place preference observed in KD animals were unexpected. Based on the reported role of opioid signaling in the “liking” component of reward, we anticipated a decrease in sweet taste preference or possibly reduced consumption of sweetened water (Castro and Berridge, 2014; Peciña and Berridge, 2005). We also hypothesized that the loss of mu and delta receptors would affect the rewarding effects of social contact, but we found no evidence of any social impairments. We were uncertain whether knockdown could affect motivation (i.e., delay discounting) or the rewarding effects of morphine, and found them to be unaltered. The simplest explanation for the normal phenotypes is that knockdown allowed sufficient opioid receptor activity to sustain reward-driven behaviors. Nevertheless, we find this possibility unlikely, as behavioral changes have been reported after antagonist treatment, which is unlikely to completely block receptor function. Alternatively, it can be argued that opioid receptors in other types of striatal neurons play the primary and essential role in reward-driven behaviors. However, this may be inconsistent with the reported role of D1-expressing neurons in mediating the reinforcing effects of rewards (e.g. Calipari et al., 2016; Cole et al., 2018; Kravitz et al., 2012). Moreover, this assumption might be counterintuitive in the context of the reported rescue of morphine-conditioned place preference in mice with *Oprm1* gene KO by the reintroduction of mu receptors in *Pdyn*-expressing (i.e., mainly direct pathway medium spiny neurons in the nucleus accumbens) cells (Cui et al., 2014) or the reported involvement of delta opioid receptors on D1-expressing neurons in resilience to social stress (Nam et al., 2019). One way to reconcile these reports and the observed phenotypes is to assume redundancy in the roles played by opioid signaling in different types of medium spiny neurons and through presynaptic receptors. Thus, opioid receptors on dopamine neurons lacking D1 receptors or on presynaptic terminals could possibly compensate for the effects of knockdown. Opioid receptors have primarily inhibitory effects on transmission, and their activation in any part of the network that drives activity, such as delta receptors on excitatory neurons in the cortex or mu and delta receptors on cholinergic neurons, could have similar effects. This is speculative and immediately raises the question of why redundancy in opioid system functions could be necessary.

### Conclusions

We found that mu and delta receptors are ubiquitously present in D1-expressing medium spiny neurons but are not essential for the control of signaling underlying reward-driven behaviors.

## Literature

Ambrose, L.M., Gallagher, S.M., Unterwald, E.M., Van Bockstaele, E.J., 2006. Dopamine-D1 and delta-opioid receptors co-exist in rat striatal neurons. Neurosci. Lett. 399, 191–196.

Bals-Kubik, R., Ableitner, A., Herz, A., Shippenberg, T.S., 1993. Neuroanatomical sites mediating the motivational effects of opioids as mapped by the conditioned place preference paradigm in rats. J. Pharmacol. Exp. Ther. 264, 489–495.

Banghart, M.R., Neufeld, S.Q., Wong, N.C., Sabatini, B.L., 2015. Enkephalin Disinhibits Mu Opioid Receptor-Rich Striatal Patches via Delta Opioid Receptors. Neuron 88, 1227–1239.

Berridge, K.C., Robinson, T.E., Aldridge, J.W., 2009. Dissecting components of reward: ‘liking’, ‘wanting’, and learning. Curr. Opin. Pharmacol., Neurosciences 9, 65–73.

Calipari, E.S., Bagot, R.C., Purushothaman, I., Davidson, T.J., Yorgason, J.T., Peña, C.J., Walker, D.M., Pirpinias, S.T., Guise, K.G., Ramakrishnan, C., Deisseroth, K., Nestler, E.J., 2016. In vivo imaging identifies temporal signature of D1 and D2 medium spiny neurons in cocaine reward. Proc. Natl. Acad. Sci. 113, 2726–2731.

Castro, D.C., Berridge, K.C., 2014. Opioid Hedonic Hotspot in Nucleus Accumbens Shell: Mu, Delta, and Kappa Maps for Enhancement of Sweetness “Liking” and “Wanting.” J. Neurosci. 34, 4239–4250.

Charbogne, P., Gardon, O., Martín-García, E., Keyworth, H.L., Matsui, A., Mechling, A.E., Bienert, T., Nasseef, T., Robé, A., Moquin, L., Darcq, E., Ben Hamida, S., Robledo, P., Matifas, A., Befort, K., Gavériaux-Ruff, C., Harsan, L.-A., von Elverfeldt, D., Hennig, J., Gratton, A., Kitchen, I., Bailey, A., Alvarez, V.A., Maldonado, R., Kieffer, B.L., 2017. Mu Opioid Receptors in Gamma-Aminobutyric Acidergic Forebrain Neurons Moderate Motivation for Heroin and Palatable Food. Biol. Psychiatry, Obesity and Food Addiction 81, 778–788.

Cole, S.L., Robinson, M.J.F., Berridge, K.C., 2018. Optogenetic self-stimulation in the nucleus accumbens: D1 reward versus D2 ambivalence. PLOS ONE 13, e0207694.

Crittenden, J.R., Graybiel, A.M., 2011. Basal Ganglia Disorders Associated with Imbalances in the Striatal Striosome and Matrix Compartments. Front. Neuroanat. 5.

Cui, Y., Ostlund, S.B., James, A.S., Park, C.S., Ge, W., Roberts, K.W., Mittal, N., Murphy, N.P., Cepeda, C., Kieffer, B.L., Levine, M.S., Jentsch, J.D., Walwyn, W.M., Sun, Y.E., Evans, C.J., Maidment, N.T., Yang, X.W., 2014. Targeted expression of μ-opioid receptors in a subset of striatal direct-pathway neurons restores opiate reward. Nat. Neurosci. 17, 254–261.

Dölen, G., Darvishzadeh, A., Huang, K.W., Malenka, R.C., 2013. Social reward requires coordinated activity of accumbens oxytocin and 5HT. Nature 501, 179–184.

Elghaba, R., Bracci, E., 2017. Dichotomous Effects of Mu Opioid Receptor Activation on Striatal Low-Threshold Spike Interneurons. Front. Cell. Neurosci. 11, 385.

Fields, H.L., Margolis, E.B., 2015. Understanding opioid reward. Trends Neurosci. 38, 217–225.

Filliol, D., Ghozland, S., Chluba, J., Martin, M., Matthes, H.W., Simonin, F., Befort, K., Gavériaux-Ruff, C., Dierich, A., LeMeur, M., Valverde, O., Maldonado, R., Kieffer, B.L., 2000. Mice deficient for delta- and mu-opioid receptors exhibit opposing alterations of emotional responses. Nat. Genet. 25, 195–200.

Fremeau, R.T., Duncan, G.E., Fornaretto, M.G., Dearry, A., Gingrich, J.A., Breese, G.R., Caron, M.G., 1991. Localization of D1 dopamine receptor mRNA in brain supports a role in cognitive, affective, and neuroendocrine aspects of dopaminergic neurotransmission. Proc. Natl. Acad. Sci. 88, 3772–3776.

Friard, O., Gamba, M., 2016,. BORIS: a free, versatile open-source event-logging software for video/audio coding and live observations. Methods in Ecology and Evolution. 7, 1325–1330.

Gangarossa, G., Espallergues, J., De Kerchove D’Exaerde, A., El Mestikawy, S., Gerfen, C., Hervé, D., Girault, J.-A., Valjent, E., 2013. Distribution and compartmental organization of GABAergic medium-sized spiny neurons in the mouse nucleus accumbens. Front. Neural Circuits 7.

Gendron, L., Cahill, C.M., Zastrow, M. von, Schiller, P.W., Pineyro, G., 2016. Molecular Pharmacology of δ-Opioid Receptors. Pharmacol. Rev. 68, 631–700.

Hamida, S.B., Boulos, L.-J., McNicholas, M., Charbogne, P., Kieffer, B.L., 2019. Mu opioid receptors in GABAergic neurons of the forebrain promote alcohol reward and drinking. Addict. Biol. 24, 28–39.

Jastrzębska, K., Walczak, M., Cieślak, P.E., Szumiec, Ł., Turbasa, M., Engblom, D., Błasiak, T., Rodriguez Parkitna, J., 2016. Loss of NMDA receptors in dopamine neurons leads to the development of affective disorder-like symptoms in mice. Sci. Rep. 6, 37171.

Kravitz, A.V., Tye, L.D., Kreitzer, A.C., 2012. Distinct roles for direct and indirect pathway striatal neurons in reinforcement. Nat. Neurosci. 15, 816–818.

Kupchik, Y.M., Brown, R.M., Heinsbroek, J.A., Lobo, M.K., Schwartz, D.J., Kalivas, P.W., 2015. Coding the direct/indirect pathways by D1 and D2 receptors is not valid for accumbens projections. Nat. Neurosci. 18, 1230–1232.

Le Merrer, J., Becker, J.A.J., Befort, K., Kieffer, B.L., 2009. Reward Processing by the Opioid System in the Brain. Physiol. Rev. 89, 1379–1412.

Ma, Y.-Y., Cepeda, C., Chatta, P., Franklin, L., Evans, C.J., Levine, M.S., 2012. Regional and cell-type-specific effects of DAMGO on striatal D1 and D2 dopamine receptor-expressing medium-sized spiny neurons. ASN Neuro 4.

Mansour, A., Fox, C.A., Akil, H., Watson, S.J., 1995. Opioid-receptor mRNA expression in the rat CNS: anatomical and functional implications. Trends Neurosci. 18, 22–29.

Matthes, H.W.D., Maldonado, R., Simonin, F., Valverde, O., Slowe, S., Kitchen, I., Befort, K., Dierich, A., Le Meur, M., Dollé, P., Tzavara, E., Hanoune, J., Roques, B.P., Kieffer, B.L., 1996. Loss of morphine-induced analgesia, reward effect and withdrawal symptoms in mice lacking the μ-opioid-receptor gene. Nature 383, 819–823.

Nam, H., Chandra, R., Francis, T.C., Dias, C., Cheer, J.F., Lobo, M.K., 2019. Reduced nucleus accumbens enkephalins underlie vulnerability to social defeat stress. Neuropsychopharmacology 44, 1876–1885.

Novak, M., Halbout, B., O’Connor, E.C., Rodriguez Parkitna, J., Su, T., Chai, M., Crombag, H.S., Bilbao, A., Spanagel, R., Stephens, D.N., Schütz, G., Engblom, D., 2010. Incentive Learning Underlying Cocaine-Seeking Requires mGluR5 Receptors Located on Dopamine D1 Receptor-Expressing Neurons. J. Neurosci. 30, 11973–11982.

Oude Ophuis, R.J.A., Boender, A.J., van Rozen, A.J., Adan, R.A.H., 2014. Cannabinoid, melanocortin and opioid receptor expression on DRD1 and DRD2 subpopulations in rat striatum. Front. Neuroanat. 8.

Panksepp, J.B., Lahvis, G.P., 2007. Social reward among juvenile mice. Genes Brain Behav. 6, 661–671.

Peciña, S., Berridge, K.C., 2005. Hedonic Hot Spot in Nucleus Accumbens Shell: Where Do μ-Opioids Cause Increased Hedonic Impact of Sweetness? J. Neurosci. 25, 11777–11786.

Perrine, S.A., Hoshaw, B.A., Unterwald, E.M., 2006. Delta opioid receptor ligands modulate anxiety-like behaviors in the rat. Br. J. Pharmacol. 147, 864–872.

Svingos, A.L., Moriwaki, A., Wang, J.B., Uhl, G.R., Pickel, V.M., 1996. Ultrastructural Immunocytochemical Localization of μ-Opioid Receptors in Rat Nucleus Accumbens: Extrasynaptic Plasmalemmal Distribution and Association with Leu5-Enkephalin. J. Neurosci. 16, 4162–4173.

Trezza, V., Damsteegt, R., Achterberg, E.J.M., Vanderschuren, L.J.M.J., 2011. Nucleus Accumbens μ-Opioid Receptors Mediate Social Reward. J. Neurosci. 31, 6362–6370.

Williams, J.T., Ingram, S.L., Henderson, G., Chavkin, C., Zastrow, M. von, Schulz, S., Koch, T., Evans, C.J., Christie, M.J., 2013. Regulation of μ-Opioid Receptors: Desensitization, Phosphorylation, Internalization, and Tolerance. Pharmacol. Rev. 65, 223–254.

Wise, R.A., 2004. Dopamine, learning and motivation. Nat. Rev. Neurosci. 5, 483–494.

Zeisel, A., Hochgerner, H., Lönnerberg, P., Johnsson, A., Memic, F., van der Zwan, J., Häring, M., Braun, E., Borm, L.E., La Manno, G., Codeluppi, S., Furlan, A., Lee, K., Skene, N., Harris, K.D., Hjerling-Leffler, J., Arenas, E., Ernfors, P., Marklund, U., Linnarsson, S., 2018. Molecular Architecture of the Mouse Nervous System. Cell 174, 999–1014.e22.

